# Culture of human retinal explants as an ex vivo model for retinal gene therapy

**DOI:** 10.1101/2024.05.06.592769

**Authors:** Almaqdad Alsalloum, Madina Khubetsova, Olga Mityaeva, Pavel Volchkov

## Abstract

Optogenetic gene therapy may employ recombinant adeno-associated virus (AAV) vectors for specific preclinical applications in the treatment of inherited retinal diseases. Human retinal explants as an ex vivo model assist determine cellular tropism and the level of subsequent gene expression followed by delivery of viral vectors. Extracts of retinal tissue can be obtained and cultured for a certain time; thus, it is possible to transduce and confirm the effectiveness of the virus using various methods, including immunohistochemical staining. Consequently, we described method for harvesting and culturing human retinal fragments from divergent parts of the retina that differ anatomically and molecularly. The fragments were fixed, sectioned and stained. A dissociation method has also been described to obtain a retinal cell suspension that can be used in other experiments such as flow cytometry or mRNA/protein level expression assays.

## 1. Introduction

Optogenetic gene therapy has been presented as a suggestion to remediate the progression of inherited retinal diseases, previously obstinate blindness-related diseases, and even to restore various types of retinal thinning and degradation [1,2]. Adeno-associated virus (rAAV) vectors for the treatment of retinal diseases have demonstrated safety, efficacy, and an absence of side effects in primary clinical trials conducted by transducing patients with RPE65 gene mutation associated with Leber’s congenital amaurosis (LCA2). In addition, improvements in visual field and function have been evaluated in these patients [3,4].

Animal and cell models have been, and still are, considered the cornerstone for evaluating preclinical AAV-related therapy, despite its lengthy, exorbitant, and indecisive results. The boundless number of AAV designs has made it possible to try-out the tropism and efficacy of AAV in wide range of animal and cell models, however, there is a discrepancy between the expression of transgenes in the organs of animal-models and human subjects [5]. The feasibility of optogenetic gene therapy accrues from the outstanding possessions of the retina, chiefly the unique immune system, the ceiling of the passive prevalence of viral vectors through its own specific barriers, and the feasibility of local injections. On top of that, restoration of visual function following photoreceptor degeneration can be achieved by effortlessly assessing using impartial and subjective functional tests [6,7]. Jacobson et al. performed authenticated information during LCA2 treatment trials on virus dispensation, toxicity, vector efficacy for targeted cell types, and novel viral vectors in the non-human primates. However, non-human primate therapeutic evaluation processing is not imperative for the final therapeutic appraisal [8].

Human retinal explants can provide details regarding viral tropism and concerning comparative efficacy [9]. Ex vivo culturing of retinal explants can be obtained in the laboratory with the restoration of functional and morphological features, despite the ineptitude in safety and biodistribution assessments [10,11]. Human retinal explants are a slightly flawed model depending on the conditions of its extraction, for instance, patients with critical retinal detachment, diabetic eye disease, or vitreoretinopathy [12]. Nevertheless, human retinal tissue is being considered as an eventual preclinical pattern and a target tissue at issue [13]. Johnson and Martin described their own system for culturing rat retinal explants for stem cell therapy [13]. Another study is based on Johnson and Martin system in which human retinal fragments were cultured in a Neurobasal-A based medium after retinal piece extraction at the time of surgery. AAV vectors were constructed and then introduced into retinal fragments with reporter genes to visualize successful transduction of retinal cells using fluorescence microscopy. In addition, post-culturing period experiments including fixation, sectioning for immunohistochemistry, or dissociation for processing mRNA/protein expression analysis were obtained [14]. Several basic conditions must be taken into account when extracting and culturing retina, including removal of the anterior part of the eye, the presence of a CO2-independent environment, the careful separation of retinal tissue from the vitreous, and the placement of retinal fragments on a Transwell membrane with a certain pore size [9]. Consequently, we attempted to extract post-mortem human retinas from patients’ eyes within a specified time frame of 6 hours as maximum to evaluate the tropism and efficacy of viral vectors on retinal cell types. In addition, we obtained a dissociation protocol to prepare cells for analysis of mRNA/proteins expression and sectioning of retinal fragments to stain specific retinal cells to determine the affinity of the virus for cell types.

## 2. Materials

Handling of the human ocular organ should be done in a clean, sterile and barren environment in a culture hood, which utilizes a positive pressure. Prepared culture media can be pre-aliquoted and stored at -20°C until use. All surgical and manipulation instruments must be sterilized, especially the retinal fragments dissection tools.

### 2.1 Dissecting procedure and explant culture

1. Hibernate-A medium (see Note 1).
2. Retinal culture medium (RCM): 500 ml DMEM/F12 (or GMEM) with 5 ml 100x glutamax or L-glutamine, 5 ml 100x non-essential amino acids, 5 ml 100x sodium pyruvate, 4 µl stock β-mercaptoethanol (16 M), 10 ml 50x NS21 supplement, 1.25 ml N-acetylcysteine (0.5 mM), and 5 ml 100x lipid concentrate. Aliquot in 50 ml per tube. Thaw the tube at 4°C for several hours before use (see Note 2).
3. Cell culture inserts made of polycarbonate in 6-well/12-well plates with a pore size of 0.4 µm.
4. 10 cm diameter petri dish.
5. 3 mL Pasteur pipettes.
6. Pipettes and filter tips.
7. Humidified incubator set to 37° C with 5% CO2.
8. Positive-pressure tissue culture hood.
9. Dry ice.
10. Corneoscleral forceps (see Note 3).
11. Scissors for corneoscleral incision (see Note 4).
12. Micro-forceps and scissors.
13. Scalpel.
14. 5 or 8 mm-diameter round punch biopsy.
15. Surgical spatula.

### 2.2 Immunohistochemistry

1. 4% Paraformaldehyde (PFA) in PBS.
2. 30% Sucrose solution in PBS.
3. Optimal Cutting Temperature medium (OCT).
4. Vinyl specimen base molds (15x15x5 mm).
5. Cryotome.
6. Poly-L-lysine-coated glass slides.
7. Mounting Medium.
8. Metal block.
9. Insulated foam Container.
10. Liquid nitrogen.
11. Washing Buffer: 0.1% Triton-X, 0.1% Tween-20 in PBS. Use 100 mL PBS as final volume.
12. Blocking buffer: Dissolve in 500 mL distilled water, 50 mL goat serum (S26 liters), 1.25 mL Triton-X, 1.25 ml Tween-20, 11.25 g glycine, 5 g bovine serum albumin (BSA), 5 phosphate-buffered saline (PBS) tablets, and 0.5 g sodium citrate. Make sure the reagents are completely dissolved. Aliquot of 5 mL tubes then freeze at -20° C for later use (see Note 5).
13. Staining Buffer: To 500 mL distilled water add 1.25 mL Triton-X, 1.25 mL Tween-20, 5 g BSA and 5 PBS tablets. Aliquot 5 mL per tube and freeze at -20° C (see Note 5).

### 2.3 Dissociation

1. Retinal ganglion cell (RGC) growth medium.
2. Hanks’ balanced salt solution (HBSS) without Ca2+ Mg2+.
3. 0.22 µm filter.
4. 5 mL syringe.
5. 40 µm cell strainer.
6. Papain solution: 1.1mM EDTA, 0.3mM β-mercaptoethanol, 5.5mM cysteine-HCl and 1 mg of papain powder (Sigma-Aldrich). To 50 mL HBSS w/o Ca2+, Mg2+ add 110 µL of 0.5M EDTA, 0.043 g of L-cysteine Hydrochloride, and 1 µL of 14.2M β-mercaptoethanol. Aliquot the previous solution by 10 mL in tube, then use 1 mg of papain powder for each tube. Filter the papain solution with 0.22 µM filter (syringe/tube). Leave the tube of papain solution open at 37ºC, 5% CO2 incubator for 30 min before using on the retinal fragment (see Note 6). Prepare the papain solution fresh as required.

## 3. Methods

### 3.1 Retina explants collection and culture

The retina is removed from the donor eye after death (post-mortem). For further use, the cornea was cut, then the retina was isolated. All specimen handling should be performed under aseptic conditions in a positive pressure tissue culture hood.

1. Pour Hibernate-A medium into a 10 cm Petri dish. Place the eyecup in the Petri dish.
2. Remove fat and connective tissue from the eyecup using corneoscleral scissors.
3. Make a circular incision between the transparent cornea and the opaque sclera (through limbus) (Fig 1a, 2a).
4. Remove the anterior segment and lens. Start removing as much as possible from the vitreous body (see Note 7).
5. Divide the eyecup into four sides to create a flower-like structure with four petals (Fig 1b, 2b).
6. The flower-structure will be clearer when the eyecup is flattened.
7. Use a 5 mm round hole punch biopsy tool for diacritic explants (Fig 1c, 2b). Press down using the punch and gently roll until an incision is made around the macula, mid-periphery, or peripheral retina. Culture each type of retina separately (see Note 8).
8. Using a Micro-forceps, try to carefully separate the retina from its RPE, which will be stuck with choroid.
9. After separation, transfer the retinal explant and the choroid-RPE tissue into transmembrane well (Fig 1d) inserted in 6 well plate individually (see Note 9).
10. Add every 3-4 pieces of the retinal explants separately per well. Make sure there is enough space between them.
11. Add 1-1.2 ml of retinal culture medium (RCM) to the wells around the transmembrane wells (see Note 10).
12. Place the 6-well plate with the explants in a humidified culture incubator set to 37°C with 5% CO2.
13. Change the medium every day by removing the day-before retinal culture medium and add new one. Applying the viral vectors can be performed after 24 hours of culturing (see Note 11).

**Fig. 1.**
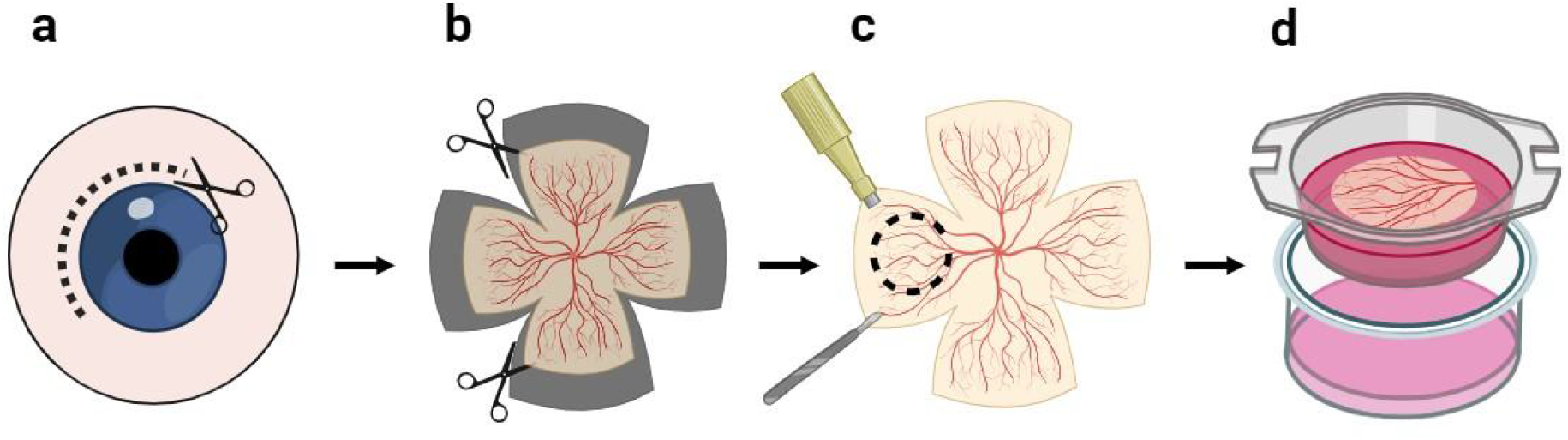
Simplified scheme for harvesting and culturing retinal explants. (**a**) Obtaining a circular incision between the transparent cornea and opaque sclera, then removing the anterior segment, lens, and vitreous. (**b**) Dividing the remaining eyecup into four center-connected pieces to create a flower-like structure with four petals. (**c**) Diacritic of retinal explants using a 5-mm round punch. Therefore, the explant can be separated from the RPE-choroid structure. (**d**) Transferring of the retinal explant into culture inserts in a well plate followed by culturing retinal tissue for use in gene therapy assessment.

**Fig. 2.**
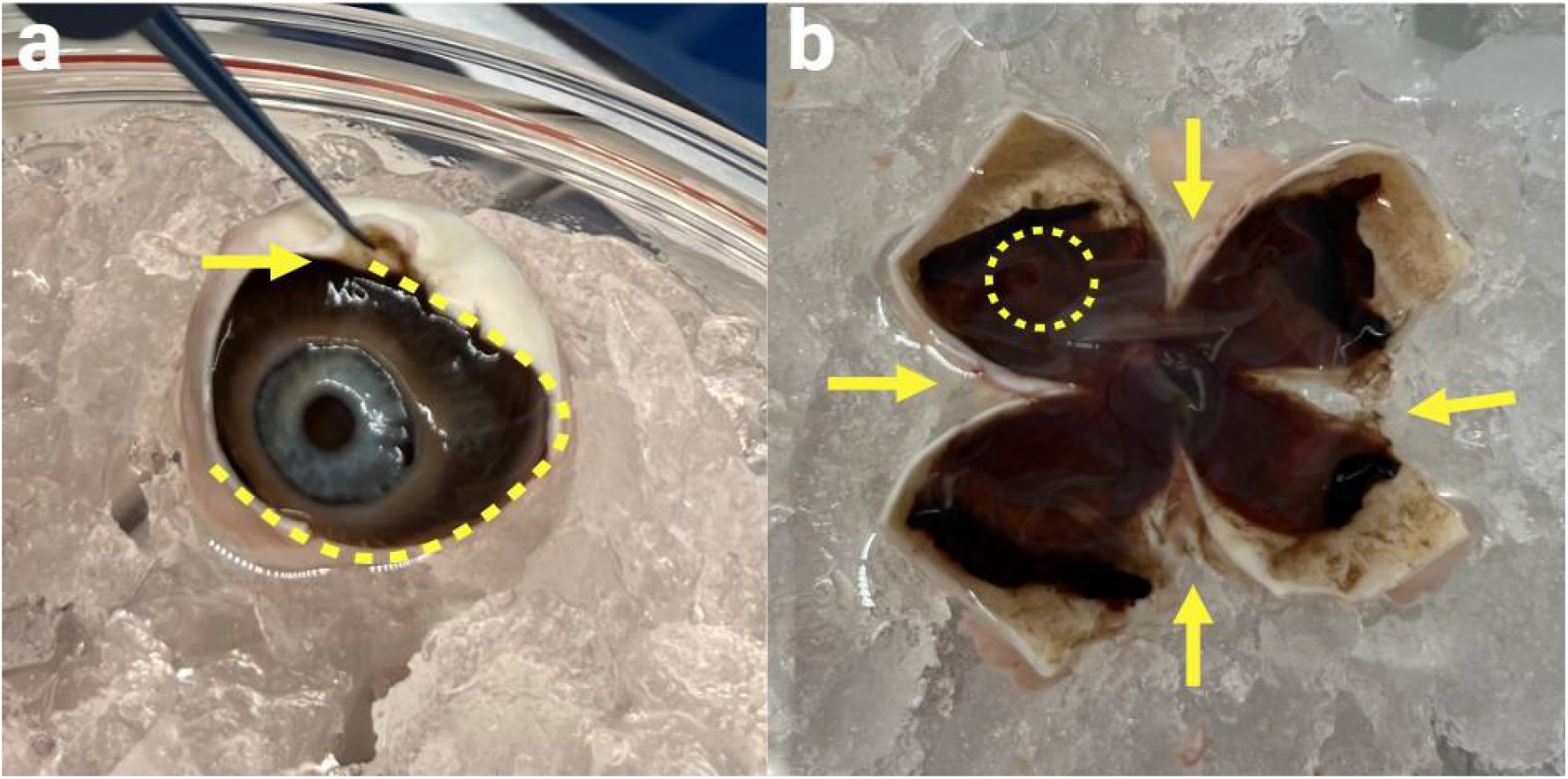
Retinal dissection and explant culture. (**a**) Human postmortem eyecup after the elimination of fat and connective tissue. A district incision is performed around the limbus of eye (yellow arrow), followed by enucleate the anterior segment and lens. (**b**) The remaining eyecup was quartered (yellow arrows) and flattened to create a flower-like structure with four petals. Therefore, a round punch was used to mark the retinal fragments (dashed line).

### 3.2 Retina transduction

1. The day after culture, it is time to transduce the retinal fragments with AAV viral vectors. Remove the medium from the well around the culture inserts.
2. Add the required viral particles to the desired part of the retina (macula, mid-periphery, peripheral retina, or RPE-choroid tissue). Use 1010 viral particles per 5 mm retinal explant fragment (see Note 12).
3. After virus injection, add 1-1.2 L of retinal culture medium around the culture inserts.
4. Return the explants back to the incubator at 37° C with 5% CO2.
5. Change the medium every day. On the 8th day after transduction, check the fluorescence of the virus (if the viral construction contains a fluorescence reporter gene). Use the explants for the required indications; immunohistochemistry or retinal tissue dissociation (see Note 13).

### 3.3 Immunohistochemical staining

1. After day 8 of transduction (see Note 14), transfer the retinal or RPE-choroid structure into a tube (see Note 15), then wash it with PBS.
2. Carefully aspirate PBS, then add buffered 4% PFA to the retinal fragments. Incubate overnight at 4° C (see Note 16).
3. Remove PFA. Then, wash 3 times with PBS.
4. Place tissue in 15% sucrose. Incubate overnight at 4° C (see Note 17).
5. Change the solution to 30% sucrose (see Note 17), and store at 4° C for one or two days (see Note 18).
6. Place the metal block into the foam container. Fill with liquid nitrogen so that the block is not completely covered. Wait until the block freezes.
7. Prepare a vinyl specimen mold tray filled with a small-scale amount of OCT.
8. Wash the sample once in PBS. Transfer the retinal fragment to separate vinyl trays (see Note 19). Cover the top with medium, do this swiftly so that the sample does not have time to reach the bottom of the tray (see Note 20).
9. Place the mold tray on the frozen metal block in the container and incubate until the matrix medium is completely frozen (see Note 21). Trays should be stored at −80° C prior to sectioning.
10. Mark the edges of the frozen OCT to indicate the location of the specimen.
11. Remove frozen OCT from the plastic mold. Fix the frozen OCT by adding more OCT to the metal platform and along the thin edge.
12. Cut on a cryotome 13 µm thick. Sections should be obtained on glass slides coated with poly-L-lysine (see Note 22).
13. For immunohistochemical staining. A standard protocol can be followed; Remove the OCT by washing with PBS for 10 minutes.
14. Remove excess PBS. Add 200 µl of blocking buffer throughout the slide. Incubate for more than 2 hours at ambient temperature.
15. Add the primary antibody at the appropriate dilution to 150 µl of staining buffer.
16. Incubate overnight at 4° C in a humid plastic chamber.
17. Remove the day-before medium, wash 2 times for 10 minutes with wash buffer.
18. Remove wash buffer, then add secondary antibody at the appropriate dilution to 150 µl of staining buffer.
19. Incubate for 4 hours at ambient temperature in the dark in a humid plastic chamber.
20. Wash as in step 17.
21. Add 150 µl of DAPI dye. Incubate for 7 minutes at room temperature in the dark.
22. Wash the sample for 10 minutes with PBS.
23. Let the sample dry and cover with mounting medium.
24. Place the cover slip on the glass slide. Allow the mounting medium to cure at room temperature before imaging (Fig 3).

**Fig. 3.**
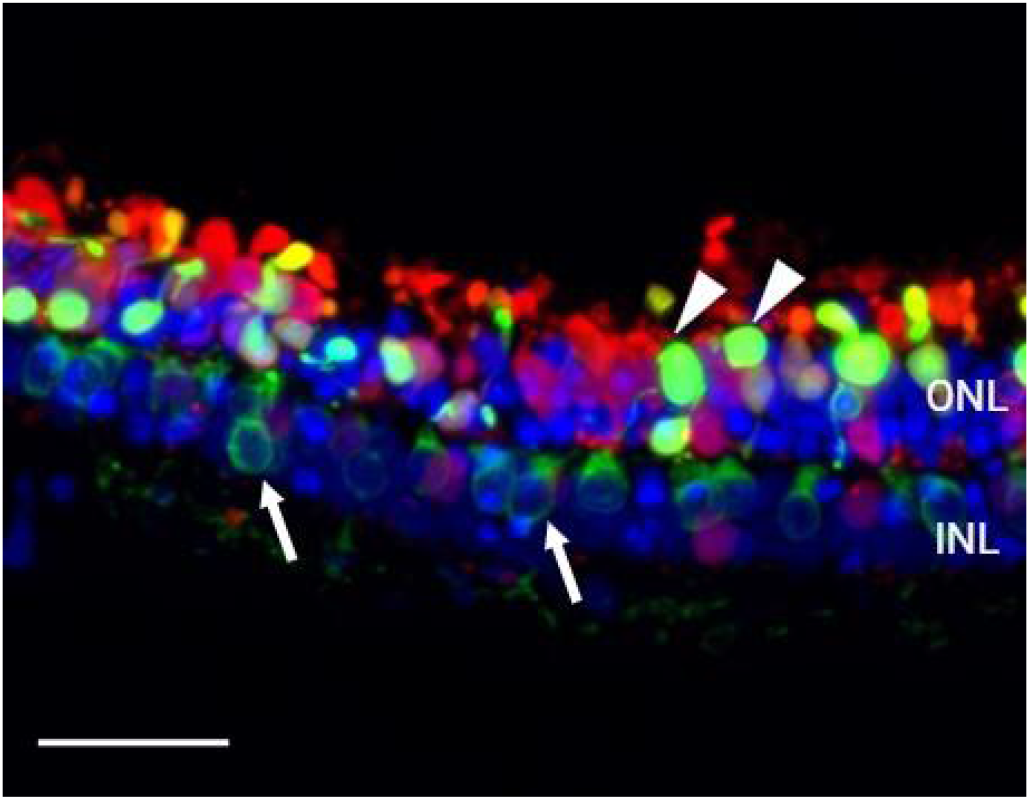
Transduction of a human retinal explant using the AAV8.GFP viral vector. Confocal microscopy image of retinal explant section after AAV8.GFP transduction, fixation, sectioning and staining. GFP expression (green) as a marker for transduction of retinal photoreceptors (arrows) and retinal bipolar cells (arrows). Recoverin as a marker of retinal photoreceptors (red). A DAPI fluorescent dye (blue) has been added to indicate the nucleus, scale bar 50 µm. INL: inner nuclear layer; ONL: Outer nuclear layer.

### 3.4 Retinal explant dissociation

1. Collect the retinal explant in a 15 ml tube.
2. Settle by gravity for 2 minutes at approximately 1600-1800 rpm, at ambient temperature to collect retinal tissue.
3. Carefully aspirate the suspension (medium) without losing retinal tissue.
4. Add 8 mL HBSS without Ca2+, Mg2+ to the retinal explant precipitate.
5. Centrifuge 7 minutes at approximately 1600-1800 rpm.
6. Discard the supernatant and resuspend the pellet in papain solution (see Note 23).
7. Incubate retinal tissue with papain solution for 20 minutes in an incubator (37°C) and shake every 4 minutes.
8. Gently resuspend the retina with papain solution more than 10 times to mechanically disengage the aggregates (see Note 24).
9. Collect the cell suspension and pass it through a 40-µm cell strainer into a new tube.
10. Centrifuge the sample at approximately 1600-1800 rpm for 5-7 minutes.
11. Discard the supernatant, then resuspend the pellet in retinal ganglion cell (RGC) growth medium for culturing (see Note 25).
12. Calculate the number of retinal cells in suspension and evaluate cell viability.
13. Use the cell suspension as a sample for mRNA/protein expression experiments. Cells can be fixed and stained for other studies such as flow cytometry analysis.

## 4. Notes

1. It is recommended to use Hibernate-A after the addition of B-27® or 100x L-Glutamine, then it can preserve healthy nerve tissue for up to 1 month at 2°C to 8°C.
2. For better sterilization and avoidance of contamination, it is recommended to add an antibiotic to the prepared medium.
3. Corneoscleral forceps with a diameter of 0.2 mm tooth (1x2 teeth) can be used in sclera dissecting process to successfully hold the sclera during dissection. The teeth can be smaller or larger, and the overall length of the forceps can be about 110 mm for easier handling.
4. One step ahead is the use of corneoscleral scissors with a flat handle and a total length of 110 mm. Scissors may have a curved blade to better cut the cornea around.
5. The frozen aliquot can be thawed before use and stored at 4°C for up to 2 weeks. It is important to filter the blocking and staining buffer after preparation and before freezing.
6. Measure the PH of the papain solution before and after incubation with the retinal explants.
7. In many cases, removing the entire body of the vitreous is quite difficult. The vitreous body, due to its stickiness, can be removed with a Pasteur pipette. Partially aspirate part of the vitreous with a pipette and try to gradually cut off the sucked part until approximately complete removal.
8. Instead of a round hole punch, micro-scissors can also be used to cut the retina into fragments. Try to gash so that the explants are as similar in size as possible. Folding of retinal fragments can be noticed during the formation of incisions. In these cases, it is recommended not to take pieces for culture. Furthermore, fragments with obvious vitreous impurities are best avoided.
9. Transmembrane wells (cell culture inserts) can be 6-well plate or less. Use the appropriate well size according to the experimental conditions.
10. Retinal explants receive nutrition and vital gases from the environment surrounding the pored membrane, therefore the environment must be sufficient for contact with the explants through the pores. Avoid high levels of media or bubbles under the bottom surface of the explant, which may affect the viability of the explant and the performance of the experiments.
11. Culture inserts should be moved with sterile forceps when changing media to avoid contamination.
12. Ensure that the added viral particles have spread sufficiently over the surface of the retinal explant. The upper surface is usually photoreceptors, especially when we assume that the results of the assessment will affect the types of photoreceptor cells. An alternative protocol for adding the viral genome is to add the total viral particles at three time points partially prior to day 8 of culture. Days 3, 5, and 7 after transduction are more common in this protocol.
13. Retinal explant size, capsid type, and AAV genome affect transduction efficiency. A negative control is needed to compare the virus signal and can be obtained by culturing explants without viral vectors.
14. Retinal culture can be cultured for up to 2 weeks in many cases. The fluorescence signal stabilized from the 8th day after transduction until the 20th day, in some cases the cell may begin to gradually degrade after the 8th day.
15. It is important to carefully transfer the explants. Avoid tearing or cruel handling, it is better to use a surgical spatula or a modified Pasteur pipette for transfer.
16. The fixation period and volume should be adjusted for the tissue of interest, for small and thin tissues 2-4 hours at room temperature or overnight fixation up to 4° C should be sufficient.
17. The retinal explant should be immersed in sucrose to confirm coverage of the entire sample.
18. Avoid contamination while retinal explants are stored in sucrose. It is recommended to start sectioning immediately after a day of treatment with 30% sucrose solution. For longer storage, use 0.1% sodium azide with 30% sucrose and store at 4°C. Sodium azide is poisonous.
19. Eliminate the PBS around the retinal explant after transferring in to the vinyl tray. A Pasteur pipette can be used.
20. The retinal tissue should be oriented. The sample should be laid in the middle of the OCT. Avoid direct tissue contact when repositioning the sample with the pipette tip.
21. Frozen samples can be stored at -80° C for a longer period of time.
22. The slide can be dried at room temperature during the day, then stored at -20° C.
23. Use 1 mL papain solution per retina in case of mouse retinal dissociation. It is also recommended to use 1 mL for 5 mm of retinal explant fragment.
24. Carefully resuspend the retina. Avoid excessive agitation which can cause the cells to burst.
25. Alternatively, HBSS can be used to resuspend the dissociated retinal explant pellet.

